# Fast and Memory-Efficient Dynamic Programming Approach for Large-Scale EHH-Based Selection Scans

**DOI:** 10.1101/2025.04.09.647986

**Authors:** Amatur Rahman, T. Quinn Smith, Zachary A. Szpiech

## Abstract

Haplotype-based statistics are widely used for finding genomic regions under positive selection. At the heart of many such statistics is the computation of extended haplotype homozygosity (EHH), which captures the decay of homozygosity away from a focal site. This computation, repeated for potentially millions of sites, is computationally demanding, as it involves tracking counts of unique haplotypes iteratively over long genomic distances and across many individuals. Because of these computational challenges, existing tools do not scale well when applied to large-scale population datasets, such as the 1000 Genomes Project, or the UK Biobank with 500,000 individuals. Optimizing computation becomes crucial when data sets grow large, especially when handling large sample sizes or generating training data for machine learning algorithms. https://github.com/szpiech/selscan

Here, we propose a dynamic programming algorithm that substantially improves runtime and memory usage over existing tools on both real and simulated data. On real phased data, we achieve 5-50x speedup with minimal memory footprint. Our simulations show an even more pronounced performance gap with large populations (up to 15x speedup and 46x memory reduction). EHH-based statistics designed for unphased genotypes run an order of magnitude faster, and multi-parameter support results in 20x runtime improvement. Source code and binaries are available at https://github.com/szpiech/selscan as selscan v2.1.

## 1 Introduction

Detecting regions of the genome influenced by positive selection has been a prominent focus for evolutionary biologists for decades. Identifying these regions provides important information about the genetic basis of adaptation, including, for example, lactase persistence (Ingram et al., 2009; Tishkoff et al., 2007), adaptation to high altitude (Szpiech et al., 2021; Liu et al., 2019), adaptation to cold climate (Cardona et al., 2014), and pesticide resistance (Hawkins et al., 2019; Hartmann et al., 2021). When an adaptive allele “sweeps” to high frequency, the neighboring sequence is also brought to high frequency, and this process leaves a distinct signature in the genome. High-frequency haplotypes and reduced genetic diversity around a specific locus are the hallmarks of an ongoing or recently completed sweep, as recombination and mutation have insufficient time to break up the underlying haplotypes. These patterns form the basis for designing haplotype-based selection statistics, and many summary-statistic-based approaches that capture these patterns have been developed in past decades to identify recent selection (Vatsiou et al., 2016; Hejase et al., 2020). Among them, iHS (Voight et al., 2006), nSL (Ferrer-Admetlla et al., 2014 ), XP-EHH (Sabeti et al., 2007), and XP-nSL (Szpiech et al., 2021 ), iHSL and rIHS (Zhao et al., 2024) are popular examples of statistics that leverage haplotype structure information based on patterns of Extended Haplotype Homozygosity (EHH). We call this suite of statistics EHH-based statistics.

Recently, a wave of next-generation sequencing has resulted in a significant increase in the number of sequenced genomes and associated data (Alonso-Blanco et al., 2016; Bycroft et al., 2018; Taliun et al., 2021). As these datasets grow, it is necessary to develop efficient algorithms and data structures to manage large-scale genotype datasets (Durbin, 2014; Melsted and Pritchard, 2011; Spence et al., 2023). Furthermore, machine learning (ML) approaches to detect signatures of positive selection (Lin et al., 2011; Pybus et al., 2015; Kern and Schrider, 2018; Sugden et al., 2018; Xue et al., 2021; Arnab et al., 2023; Lauterbur et al., 2023 ) are becoming popular. These approaches require large-scale training data and often use multiple summary statistics as input features to train their model. Since each summary statistic captures different genetic patterns, and is well-powered for different parts of the parameter space, combining them can help determine whether a selective sweep likely caused the observed values. As a result, these statistics need to be computed many times on many different training data with varying demographic parameters. To accomplish this at scale, computation of these summary statistics must be fast and seamless. Additionally, using the same original data set with different filtering criteria can lead to entirely different biological conclusions, making it important to reanalyze with different filtering and stopping conditions (Hemstrom et al., 2024). Therefore, these ML based methods will greatly benefit from optimizing the computation of individual statistics, as well as simultaneous computation of multiple statistics with varying parameters.

Several tools in population genetics analyses use robust data structures to exploit LD decay patterns to improve efficiency, for instance, combinatorial pattern matching for imputation and phasing (Durbin, 2014) and coalescent simulations (Kelleher et al., 2016). However, development of efficient algorithms for detecting sweeps, particularly on large-scale datasets, have been limited (although see (Maclean et al., 2015). Tools like `hapbin` (Maclean et al., 2015), `selscan` (Szpiech and Hernandez, 2014; Szpiech, 2024) and `rehh` (Gautier and Vitalis, 2012; Gautier et al., 2017) are all designed to compute iHS and XP-EHH efficiently. There are other tools that compute iHS such as `vcflib` (Garrison et al., 2022, 2021) and `sgkit` (Miles et al., 2024), but they are not specifically optimized for computation of EHH-based statistics. `rehh` is an R package to compute iHS and relevant statistics introduced around the same time as `selscan` and `hapbin`, which was later improved for performance (Gautier et al., 2017). `Hapbin` achieves efficiency by means of a multithreaded algorithm and bitwise operations and is the most computationally efficient tool. Although `hapbin` is the fastest among the existing state-of-the-art tools to compute EHH, iHS and XP-EHH, its functionalities are quite limited compared to `rehh2` and `selscan`. It does not support other EHH-based statistics nor support common file types (such as VCF, TPED, and their gzipped versions) which are supported by `selscan` and `rehh2`. Even so, when analyzing selection signatures in a high-dimensional cattle dataset (Nawaz et al., 2024) consisting of 10K samples and 7 million SNPs, `hapbin` reached its limitation, requiring alternative data pre-processing to use `hapbin`. `rehh2` and previous versions of `selscan`, on the other hand, have more functionalities and are used more widely; however, they do not scale to large datasets. Furthermore, `hapbin` does not support unphased data like `selscan v2.0` and `rehh2`. It was shown in Szpiech, 2024 that larger sample sizes are required for having good power for unphased analysis as it loses some power by dropping phase information. An efficient algorithm for unphased calculation is therefore imperative to handle larger sample sizes.

In order to address these scalability challenges for both individual large datasets and ML applications, we release a major update to `selscan` versioned as `selscan v2.1`, which removes the performance bottleneck of `selscan` by employing a faster and more memory-efficient algorithm. Furthermore, we add support to compute multiple statistics with multiple parameter settings without needing to rerun the program multiple times. In this paper, we describe the new dynamic programming algorithm and compare performance to other tools. This improved version works on massively large datasets which could not be handled by previous versions of `selscan`. We see speedups of 2x-13x over `hapbin` and 9x-34x over rehh2, along with memory improvements from 2x-46x over `hapbin` and 137x-733x over rehh2, even when limiting all tools to 48-hour run times.

## 2 Materials and Methods

In this section, we describe the core components and innovations behind the development of selscan v2.1, a re-engineered and optimized tool for computing EHH-based selection statistics. selscan v2.1 introduces several improvements over existing tools, including a dynamic programming algorithm for speed, memory-efficient data structures, and a multi-parameter execution strategy for robust and flexible analysis. It takes as input biallelic SNP data in VCF, TPED, or HAP format. It supports both standard and transposed HAP formats, where the standard format represents haplotypes as rows and SNPs as columns, and the transposed format (e.g., from vcftools or shapeit) represents SNPs as rows and haplotypes as columns. Information about genetic and physical distance is provided either through a separate MAP file or embedded within TPED or VCF files. It outputs one or more selection statistics based on user-defined parameters.

Section 2.1 defines the EHH-based statistics and their mathematical formulations. Section 2.2 introduces a dynamic programming framework for efficient computation. Section 2.3 describes our bitset-based storage optimization. Section 2.4 outlines our multi-parameter execution strategy. Finally, Section 2.5 details the datasets and experimental setup used to evaluate performance.

### 2.1 EHH-based Statistics

#### 2.1.1 Input and Output

Given *K* binary sequences representing a sample of *K* chromosomes, each of length *N*, we define a *haplotype matrix* **H** ∈ {0, 1} ^*K×N*^, where each of the *K* rows corresponds to a haplotype and each of the *N* columns corresponds to a bi-allelic variable genetic site (i.e., a marker locus). Alternatively, this matrix can be viewed as a list of *K* binary strings of length *N*, each representing a haplotype. In this encoding, ancestral alleles are denoted by 0 and derived alleles by 1.

Our algorithm takes a haplotype matrix **H**(*K, N* ) as input for single-population statistics such as iHS, nSL, iHH12, EHH, and soft-EHH. For cross-population statistics (e.g., XP-EHH and XP-nSL), it requires two such matrices, **H**_1_(*K, N* ) and **H**_2_(*K, N* ), representing two populations. Additionally, we require auxiliary vectors of physical or genetic coordinates of length *N*, corresponding to the columns of the haplotype matrix, to associate each locus with its genomic position. In all cases, the selection statistic is computed independently for each of the *N* loci.

#### 2.1.2 Modeling Input Haplotypes as Multisets

In order to define extended haplotype homozygosity (EHH) based statistics precisely and compute it efficiently, we represent a haplotype column slice as a *multiset* of haplotype substrings. A multiset allows repeated elements, capturing both the identity and frequency of haplotypes without regard to their ordering, which is irrelevant for EHH computation. This abstraction simplifies definitions and algorithmic operations, as EHH depends only on the distribution of haplotype patterns, not their positions in the matrix.

By modeling haplotypes as multisets, we reduce unnecessary complexity and focus on the core elements that influence EHH and related statistics.

#### 2.1.3 Formal Definition of EHH in Both Directions

Here we define Extended Haplotype Homozygosity (EHH), as the main algorithmic contribution of this paper is captured in implementation of EHH. The other statistics listed above are all functions of the core EHH computations.

Consider a focal locus *i* (0 ≤ *i < N* ) in a binary haplotype matrix **H** ∈ {0, 1} ^*K×N*^, where *K* is the number of haplotypes and *N* is the number of loci. The goal is to compute the *extended haplotype homozygosity* (EHH) from the focal locus *i* to any other locus *j* (0 *≤ j < N* ).

Let ℋ_*i,j*_ denote the multiset of substrings extracted from each haplotype (row of **H**), from locus min(*i, j*) to max(*i, j*), inclusive. That is, for each row *r*, define: *x*_*r*_ = **H**_*r*,min(*i,j*):max(*i,j*)_ and 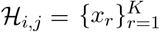. Then, the extended haplotype homozygosity from locus *i* to *j* is defined as:

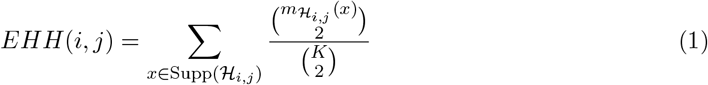

Here 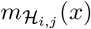 denotes the multiplicity of *x* in ℋ_*i,j*_, and Supp( ℋ_*i,j*_) denotes the set of unique substrings in ℋ_*i,j*_. This equation measures the probability that two randomly chosen haplotypes are identical between loci *i* and *j*. The definition is symmetric in *i* and *j*, and thus applies in both upstream (*j > i*) and downstream (*j < i*) directions.

##### Example

Consider the haplotype matrix **H** *∈* {0, 1}^8*×*6^:

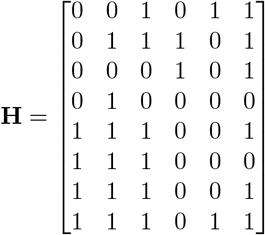

Let the focal locus be *i* = 0. We compute *EHH*(0, 3), using substrings from columns 0 to 3:

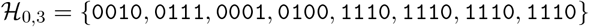

Compute multiplicities: *m*(0010) = 1, *m*(0111) = 1, *m*(0001) = 1, *m*(0100) = 1, *m*(1110) = 4. Then:

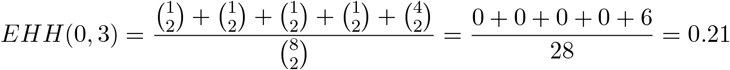

In some cases, we may wish to compute haplotype homozygosity within a subset of samples, i.e., those that carry a particular *core haplotype* at the focal locus *i*. Let 𝒞 := {0, 1} denote the set of possible alleles at locus *i*, and let *c ∈* 𝒞 be the chosen core allele. Define 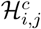 as the multiset of substrings from haplotypes that carry allele *c* at locus *i*, extending to locus *j*.

Then, the extended haplotype homozygosity among samples with core allele *c* is defined as:

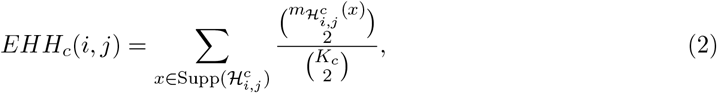

where *K*_*c*_ is the number of haplotypes carrying allele *c* at locus *i*, and ∑_C∈ *𝒞*_ *K*_*c*_ = *K*. For example, if the core locus is *i* = 1 and allele 1 is considered the derived core haplotype, then the distinct observed haplotypes extending to locus *j* = 2 are 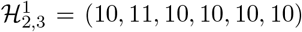. Similarly, 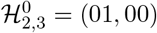 for core allele 0. Note that:

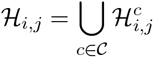

Suppose SNP 0 is chosen as the core, and samples 0 through 6 carry allele 1 at this locus. As we move upstream (rightward) from the core, over loci *upstream*(0) = {0, 1, 2, 3, 4, 5}, we observe the decay in EHH values for each core allele. The computed values are:

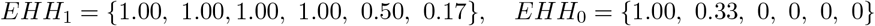

This indicates that the haplotypes carrying the derived allele (1) retain extended homozygosity over longer distances, while homozygosity among the ancestral allele (0) decays rapidly. Such patterns are indicative of recent positive selection acting on the derived allele. Figure 1 conceptually illustrates the patterns for *EHH*_1_ and *EHH*_0_ when the 1 allele is adaptive. Note that when fewer than two haplotypes are considered, EHH cannot be defined because no pairwise comparison is possible. In such cases, we set EHH to 0, since no extended haplotype homozygosity can be observed.

**Figure 1:**
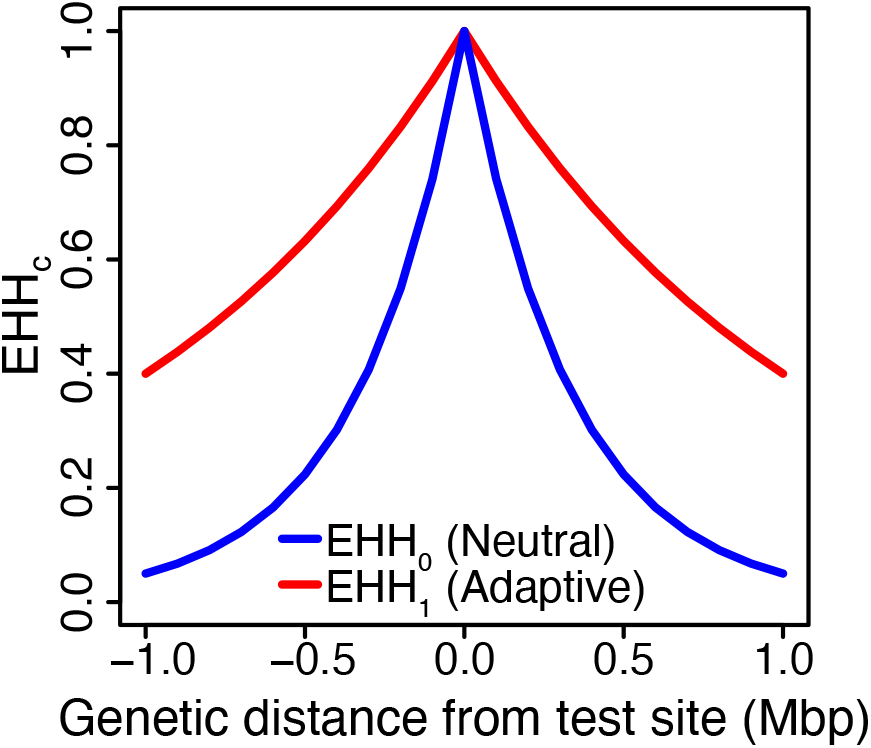
Conceptual illustration of *EHH*_*c*_. EHH is computed for a fixed core locus *i* across varying positions *j*, with upstream (right) and downstream (left) directions shown. The curves represent EHH decay for haplotypes containing the two core alleles at the focal site. The 1 allele (red) is adaptive, and therefore *EHH*_1_ decays slowly. The 0 allele (blue) is neutral, and therefore *EHH*_0_ decays quickly.

#### 2.1.4 Integrated Haplotype Score (iHS)

As established in the previous section, our goal is to track the decay of extended haplotype homozygosity (EHH) values. A favored (adaptive) allele typically maintains longer-range haplotype homozygosity, resulting in a larger area under its EHH curve. This asymmetry is quantified by the iHS statistic (Voight et al., 2006).

To compute iHS at a given site *i*, we first calculate the integrated haplotype homozygosity (iHH) for both the ancestral (*c* = 0) and derived (*c* = 1) alleles, using trapezoidal quadrature applied to the EHH curves obtained via Equation 2. Let 𝒞 := {0, 1} denote the set of core alleles. The total integrated haplotype homozygosity for allele *c* at site *i* is defined as:

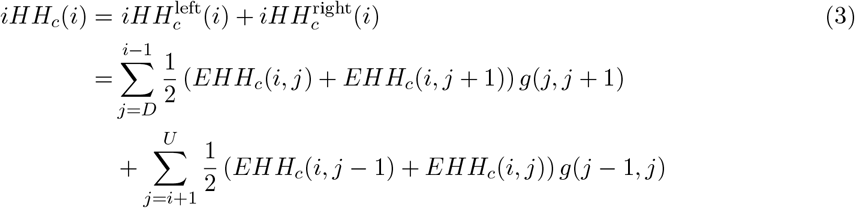

Here, *D* is the leftmost locus (downstream of *i*), and *U* is the rightmost locus (upstream of *i*) used in integration, subject to predefined distance or EHH cutoff thresholds. The function *g*(*j*^*′*^, *j*) denotes the genetic distance between loci *j*^*′*^ and *j*.

Once *iHH*_0_(*i*) and *iHH*_1_(*i*) are computed, the unstandardized iHS is calculated as:

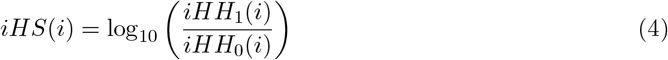

This process is repeated for each locus *i* ∈ {0, 1,. .., *N* −1} . In Figure 1, the area under each curve corresponds to the integrated haplotype homozygosity (*iHH*_1_ or *iHH*_0_). As the adaptive allele maintains longer-range haplotype homozygosity, it results in *iHH*_1_ *> iHH*_0_, and a positive iHS score.

In our previous example, consider site *i* = 0. Assume positions along the genome are evenly spaced and the spacing between consecutive positions as 1 unit. The EHH curves yield *iHH*_1_ = 1.865 and *iHH*_0_ = 0.5. Then, the iHS is:

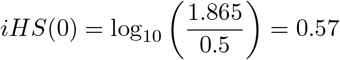

Positive iHS indicates that allele 1 has slower EHH decay, consistent with a selective sweep. In practice, this raw iHS value is calculated across all SNPs in the dataset. High-magnitude values (positive or negative) may be indicative of selection. However, to properly assess significance, iHS values are standardized within allele frequency bins to account for frequency-dependent haplotype lengths. This normalization ensures that deviations are interpreted relative to neutral expectations for alleles of similar frequency, thereby improving robustness to demographic effects.

#### 2.1.5 Cross-population Extended Haplotype Homozygosity (XP-EHH)

This idea can be extended to compare the decay of EHH patterns between two populations, which is the basis for the XP-EHH statistic (Sabeti et al., 2007).

To compute XP-EHH at a given site *i* comparing populations *A* and *B*, we first calculate the integrated haplotype homozygosity (iHH) for all haplotypes in population *A* and all haplotypes in population *B*, using trapezoidal quadrature applied to the EHH curves obtained via Eq. 1. The total integrated haplotype homozygosity for population *P* at site *i* is defined as:

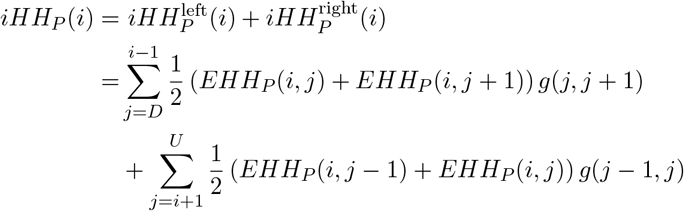

Where *EHH*_*P*_ (*i, j*) is the EHH (from Eq. 1) computed among haplotypes in population *P* . *D* is the leftmost locus (downstream of *i*), and *U* is the rightmost locus (upstream of *i*) used in integration, subject to predefined distance or EHH cutoff thresholds. The function *g*(*j*^*′*^, *j*) denotes the genetic distance between loci *j*^*′*^ and *j*.

Once *iHH*_*A*_(*i*) and *iHH*_*B*_(*i*) are computed, the unstandardized XP-EHH is calculated as:

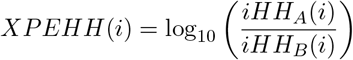

This process is repeated for each locus *i* ∈ {0, 1,…, *N* −1} . In practice, this raw XP-EHH value is calculated across all SNPs in the dataset. High-magnitude positive values may be indicative of selection in population A and high-magnitude negative values may be indicative of selection in population B. XP-EHH values are typically standardized genome-wide, although they do not have the same frequency-dependent haplotype length issue as iHS.

#### 2.1.6 Other EHH-based statistics

Although nSL (Ferrer-Admetlla et al., 2014) and XP-nSL (Szpiech et al., 2021) were not defined with respect to EHH, Ferrer-Admetlla et al. showed that nSL (and by implication XP-nSL) can be re-written in terms of EHH and iHH. In this case, the difference between nSL and iHS (and between XP-EHH and XP-nSL) is that, when calculating iHH, *g*(*j*^*′*^, *j*) denotes the number of loci between *j*^*′*^ and *j* instead of the genetic distance.

The other statistics implemented in `selscan`, iHH12 and soft-EHH, are straightforward functions of EHH, and we omit their definitions here for brevity.

### 2.2 Algorithm to Compute Phased EHH

We present an efficient algorithm for computing EHH-based statistics, formulated in terms of Eq. 1. The extension to Eq. 2 is straightforward. The scale of efficiency improvements remain unaffected by whether the algorithm is phased or unphased. Therefore, all descriptions here focus on the phased case based on Eq. 1. Unphased versions of iHS, nSL, XP-EHH, and XP-nSL (Szpiech, 2024) are valuable when phased data are unavailable, so accelerating them is equally important. We therefore implement this extension and demonstrate corresponding efficiency gains. In this setting, haplotypes originally encoded as binary 0/1 for ancestral and derived alleles are collapsed into multi-locus genotypes (MLGs) (Harris and DeGiorgio, 2020; Harris et al., 2018; DeGiorgio and Szpiech, 2022; Szpiech, 2024) with three states per locus: 0 for homozygous ancestral, 1 for heterozygous, and 2 for homozygous derived. The details of extension to unphased case for the statistics implemented in `selscan v2.1`, while not central to the efficiency results, can be found in prior work (Szpiech, 2024).

#### 2.2.1 Dynamic Programming Approach to Compute Phased EHH

##### Problem Definition

To compute EHH efficiently for a range of positions, we reparameterize using a focal locus *i* and define the length-*m* haplotypes as the multiset ℋ_*m*_ composed of substrings from position *i* to *i* + *m*. Then:

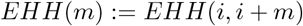

Our goal is to compute EHH for *m* = 0, 1,…, *U*, where *U < N* is an upstream cutoff. We observe that extending haplotypes from length *m−* 1 to *m* corresponds to appending a new bit (either 0 or 1) to each string in ℋ_*m−* 1_, forming ℋ_*m*_.

This allows us to define a recurrence:

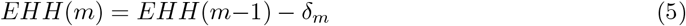

where the decrement or *drop* in homozygosity at step *m* is:

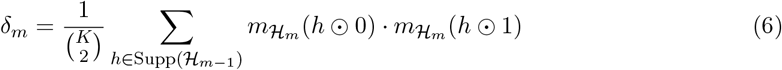

In this expression, Supp(ℋ_*m−*1_) is the set of unique prefixes of length *m−*1, and 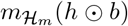 counts how many haplotypes in ℋ_*m*_ are formed by appending bit *b ∈* {0, 1} to *h*. The initial value is:

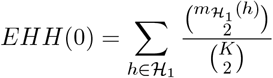

This recurrence avoids recomputing EHH from scratch for every *m*, and instead incrementally updates it by tracking changes due to newly introduced differences. Computing EHH from scratch (as in Eq. 1) would require repeatedly counting all distinct haplotypes of length *m*, which becomes increasingly costly as *m* grows and the haplotype strings become longer and more diverse. A proof of the main recurrence (Eq. 5 and Eq. 6) underlying the EHH calculation speedup is provided in the Supplementary Materials.

##### Efficient Drop Calculation via Haplotype Families

To further improve efficiency, we introduce *haplotype families*, grouping haplotypes by unique prefixes *h* ∈ ℋ _*m −* 1_. A family’s size is given by its prefix’s multiplicity, reducing the need to scan the full multiset at each step and simplifying bookkeeping. A family *splits* at position *m* if its prefix *h* extends to both *h* ⊙ 0 and *h* ⊙ 1, meaning some haplotypes continue with 0 while others continue with 1. If a prefix extends to only one of these, it does not contribute to a split.

Eq. 6 sums over all unique prefixes, regardless of whether they split. This offers conceptual clarity but incurs high computational cost. Instead, in the improved approach, we focus on *splitting families*, those that force a divergence by extending to both *h* ⊙ 0 and *h* ⊙ 1. Families extending to only one do not affect EHH and are excluded. We define the set of such *splitting families* as:

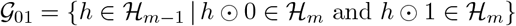

This set exclusively includes those prefixes in ℋ_*m −* 1_ that give rise to *both* 0- and 1-extended haplotypes in ℋ_*m*_, and hence *force a split* in their respective haplotype family. We refine the drop calculation as:

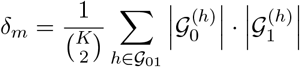

where 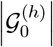 and 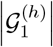 count haplotypes from prefix *h* ending in 0 and 1, respectively. This formulation improves efficiency to Θ(number of splits), which is 𝒪 (*p*_*m*_), where *p*_*m*_ is the number of 1 alleles at position *m*.

#### 2.2.2 Implementation of the dynamic programming algorithm

Our algorithm incrementally computes EHH by tracking how haplotype groups, called *families*, diverge across genomic positions. Two haplotypes belong to the same family at position *m* if their substrings from the core locus *i* to position *m* are identical. Each family is assigned a unique *color* to denote its identity.

At the core locus, haplotypes are grouped into two initial families based on whether their allele is 0 or 1. As we move rightward (i.e., increasing *m*), we check which samples carry a 1 at position

*m*. If all members of a color group (=family) have the same allele at *m*, the group (=family) stays together. If some have a 1 and others have a 0, the group (=family) *splits*: the 1-carrying members receive a new color and the others retain the original. Only these splits trigger updates to the EHH value.

This color-splitting process is tracked using compact data structures. At each position *m*, the algorithm maintains four variables: `color[ ]` (array of length *K* storing each sample’s family assignment), `count[ ]` (array counting how many haplotypes belong to each color), `numColors` (total number of distinct families so far), and `EHH` (the current extended haplotype homozygosity value). To minimize memory, only two such states are stored at a time: one for position *m* − 1 and one for *m*. This two-column design ensures constant memory usage and scales efficiently to large haplotype datasets. The key efficiency gain comes from avoiding full recomputation of EHH from scratch at every position. Instead of recounting all substrings of length *m*, we only process the families that split which is typically a small subset of all samples, thereby yielding significant speedups.

Figure 2 illustrates this process. This figure shows how haplotypes start as a single color and as the algorithm scans across positions along the genome, haplotypes split into new colors only when necessary. Updates happen only where split occurs (highlighted in bold). The overview pseudocode is provided in Algorithm 1, and detailed pseudocode runtime and memory analysis appears in the supplementary materials.

**Figure 2:**
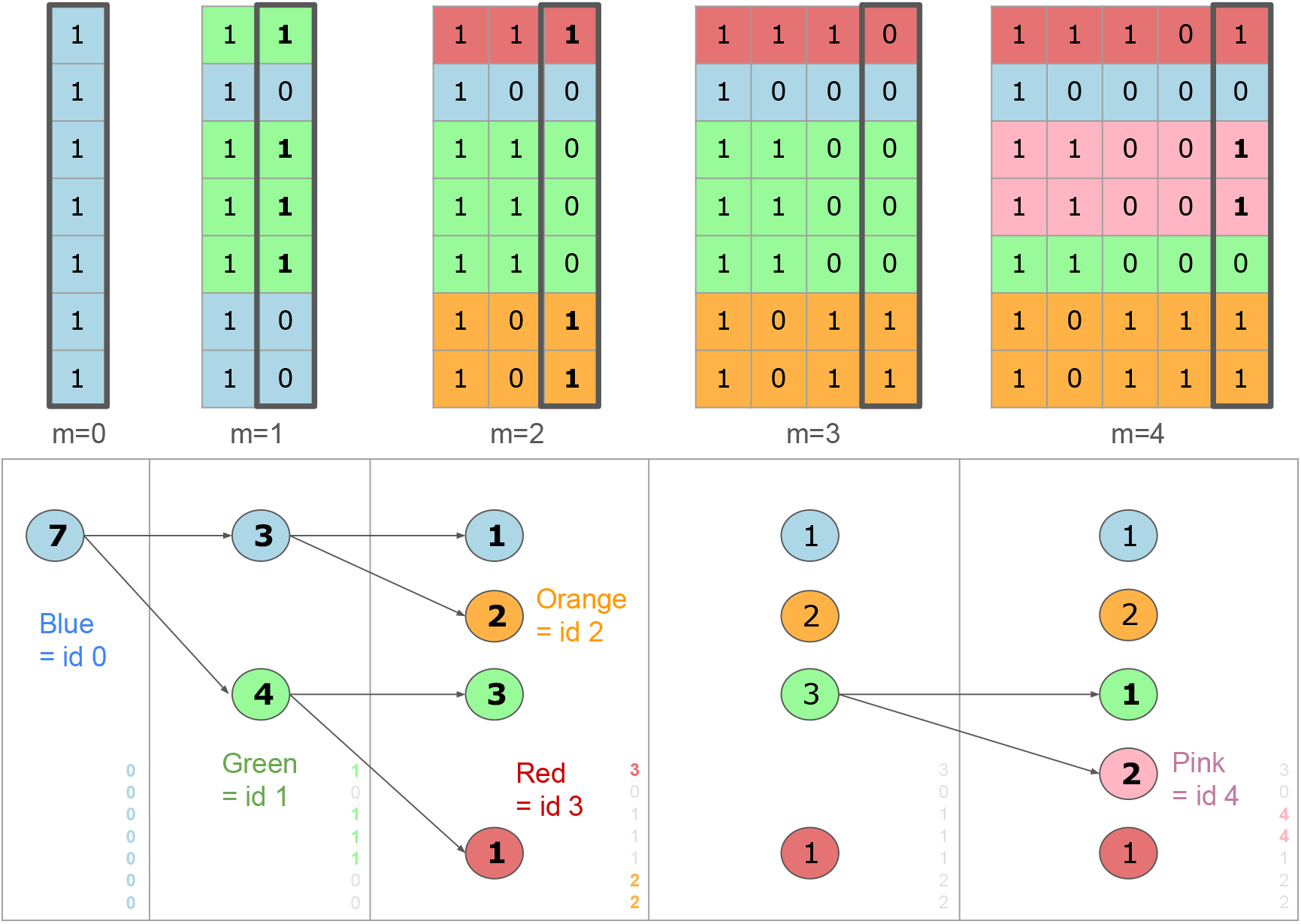
Illustration of the color-splitting algorithm on a 7 *×* 5 haplotype matrix with haplotypes (11101, 10000, 11001, 11000, 10111, 10111). The top panel shows how color assignments evolve column-wise as the algorithm progresses from left to right. The bottom panel displays the state of the core data structures in memory at each step. Each color circle represents a haplotype family, and the number inside indicates the count of samples in that family. Splits in haplotype families are shown using pairs of arrows. Note that although haplotype strings grow in length at each step, the memory state remains constant: only two columns’ worth of state variables (`count, color, numColors, EHH`) are stored at a time. The numbers in the bottom right corner represent the values of array `count`. Bold entries highlight the rows that triggered a split, corresponding to the runtime cost at that iteration. For example, at *m* = 4, color *green* splits: samples 2 and 3 carry a ‘1’ at locus 4 while sample 4 does not, resulting in a 2–1 split. A new color, *pink*, is created and paired with the original. Colors like *orange* and *red* do not split because all associated haplotypes are consistent at that locus. By tracking family splits using the prefix structure and updating EHH incrementally, the algorithm achieves linear time and low memory usage.

##### Pseudocode

The following pseudocode summarizes the core logic of the algorithm, which assumes that the focal site is not monomorphic. In practice, the case where the focal site is monomorphic (i.e., all 0s or all 1s) is handled by initializing with a single color.

###### Algorithm 1: Overview of Incremental EHH Update via Iterative Color-Splitting Algorithm

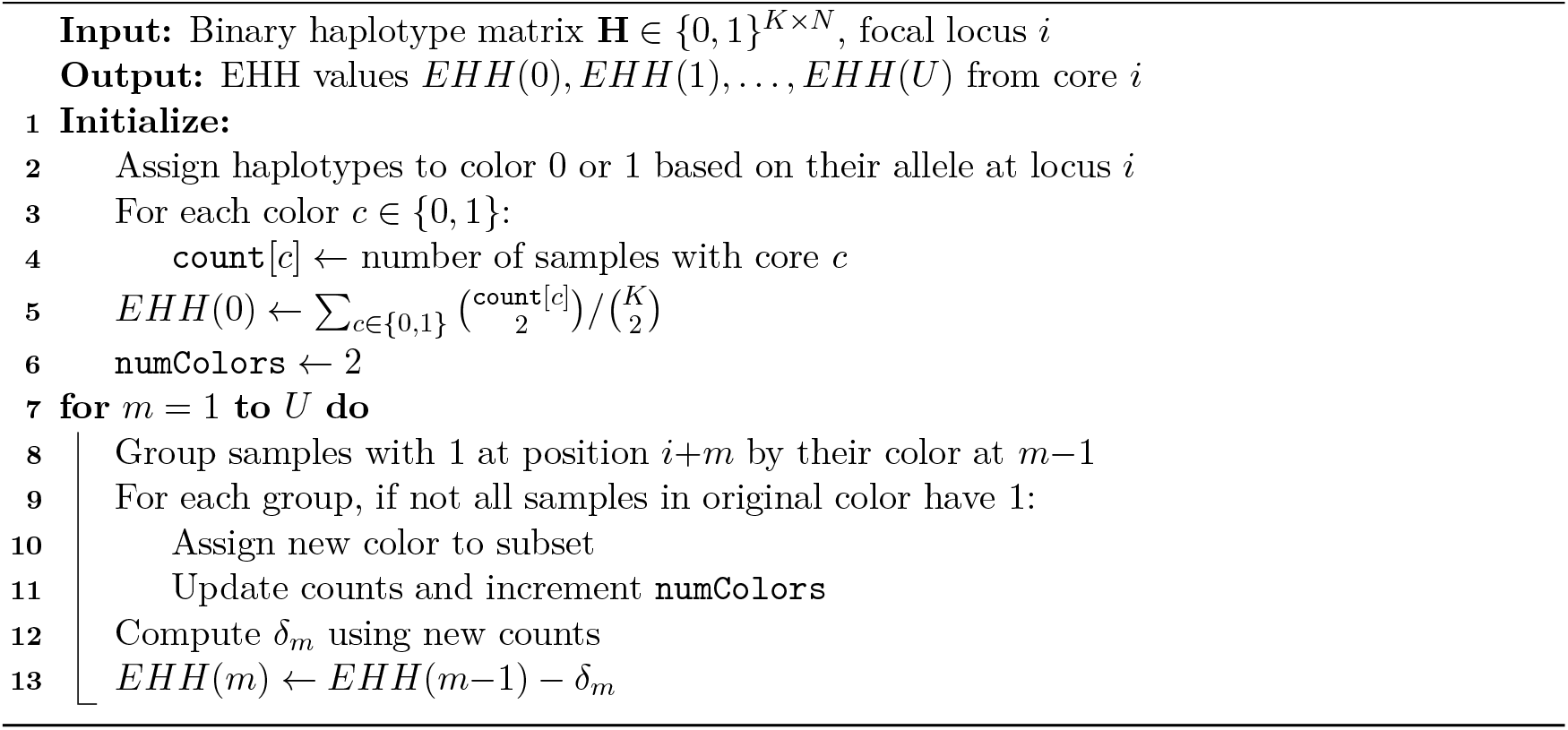

#### 2.2.3 Aggregating EHH values into summary statistics

##### From EHH to iHH

During rightward scanning, each time *EHH*(*j*) is updated, the integrated haplotype homozygosity *iHH*^right^ is incrementally accumulated as described in Eq. 3. A similar pass is made in the leftward direction to compute *iHH*^left^. After both passes, the total integrated value is computed as:

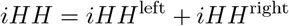

##### From iHH to summary statistics

Because EHH computation at each focal locus is independent of others, this step is highly parallelizable. As in the original `selscan`, loci are processed in parallel. However, to efficiently manage variation in runtime across loci, we employ a thread pool rather than launching separate threads per task. Once *iHH* values are computed, summary statistics such as iHS, XP-EHH, etc. are derived. This final aggregation and output step incurs negligible runtime compared to the core EHH computation.

### 2.3 Optimizing Storage of the Haplotype Matrix Using Bitsets

Selscan represents the haplotype matrix **H**(*K, N* ) as an array of *K* binary strings of length *N*, consuming one byte per character. In `selscan v2.1`, we optimize memory by storing each column (locus) as a bitset across samples, reducing the memory footprint by a factor of 8.

Let *p*_*j*_ ⊆ {0,. .., *K−*1} denote the indices of haplotypes with allele 1 at locus *j*. We store each *p*_*j*_ as a packed bit vector using *⌈K/*64*⌉* blocks of 64-bit unsigned integers (`uint64`), enabling compact storage and efficient access. For instance, in our example, we have *p*_0_ = {0, 1, 2, 3, 4, 5, 6} and *p*_3_ = {5, 6} . Here, *p*_0_ is stored as the bit pattern `11111110`, and *p*_3_ is stored as the bit pattern `00000110` (with trailing zeros for alignment).

This representation supports fast iteration over set bits (i.e., samples with allele 1 at locus *j*) using low-level hardware instructions, allowing Θ( |*p*_*j*_| ) time complexity per locus. Bit-level parallelism also improves cache locality and speeds up EHH computations.

For unphased input, we maintain two bitsets per locus: one for alternate homozygotes (both alleles are 1) and one for heterozygotes (one allele is 1). Reference homozygotes (both alleles are 0) can be inferred implicitly.

### 2.4 Boosting Efficiency via Multi-parameter Execution Strategy

Our tool accommodates multiple parameters for EHH-based statistics while ensuring runtime and memory optimizations. Users can provide variety of configurations such as minor allele frequency (MAF) thresholds, maximum physical or genetic distance from core haplotype, EHH decay cutoffs (e.g., stop when EHH < 0.05), and minimum core haplotype frequency to tailor analyses to different biological contexts. The tool also allows computation of multiple statistics such as iHS, XP-EHH, XP-nSL, and iHH12, in a single run. This multi-parameter execution strategy avoids redundant parsing of input and initialization of binary matrices, which are typically time-consuming for large genomic datasets. By reusing the loaded haplotype matrix and internal data structures across analyses, the tool minimizes I/O overhead and improves efficiency, making it especially suitable for genome-wide scans under diverse parameter settings. The details of each of these parameters are in supplementary.

### 2.5 Datasets and experimental setup

In order to test the effectiveness of the algorithm described in Section 2.2, we calculated selection statistics on a variety of real and simulated datasets. The characteristics of all the datasets we used are given in Table 1. The first three datasets for single population tests (chr19, chr10, chr6) are from 1000 genomes GRCh38 phase III project (release 20130502) (Lowy-Gallego et al., 2019). Whereas correct application of these single population selection statistics requires analyzing populations separately, here we intentionally combine samples across populations in order to evaluate a “large-sample” scenario. Datasets labeled *CEU* and *CHB* are subsets of individuals indicating two different populations and use chromosome 1. *CEU* is population code for Utah Residents with Northern and Western European Ancestry and *CHB* is for Han Chinese population in Beijing. We only kept the sites present in both populations.

**Table 1:** Description of datasets used in our experiments. (K = thousands, M = millions, B = billions). For cross-population statistics, sites were not filtered based on low MAF, hence marked as NA. For single population statistics, low MAF sites were filtered using the maf filter options in the tools benchmarked.

| Type | Dataset | n. Individuals | n. Haplotypes | n. Sites | n. Filtered Sites<br>(MAF ≥ 0.05) |
| --- | --- | --- | --- | --- | --- |
| Single<br>Population | Chr 19 | 3,202 | 6,404 | 1.5 M | 0.2 M |
|  | Chr 10 | 2,504 | 5,008 | 3.8 M | 0.35 M |
|  | Chr 6 | 3,202 | 6,404 | 4.8 M | 0.46 M |
|  | Chr 1 | 2,504 | 5,008 | 6.2 M | 0.53 M |
|  | Sim. sweep | 10,000 | 20,000 | 10 M | 12 K |
|  | Sim. neutral | 5,000 | 10,000 | 1 B | 1.2 M |
| Cross<br>Population | CEU chr1 | 99 | 198 | 6.2 M | NA |
|  | CHB chr1 | 103 | 206 |  |  |

We also performed simulation to test for scalability. We generated two simulated datasets using `msprime` (v1.2.0), one under a neutral process (mutation and recombination rates of 10^*−*8^) and one under selective sweep. For the sweep simulation, we set the simulated genome length (*L*) to 10 Mbp and the effective population size (*N*_*e*_) to 5000, with a mutation rate of 10^*−*7^ and a recombination rate of 10^*−*9^. The beneficial allele was placed at *L/*2, with its frequency increasing from 0.1 to 0.9, a selection coefficient of 0.25, and a time step of 10^*−*6^.

All of these datasets were filtered using `vcftools` and `bcftools` to retain phased, biallelic SNPs without any missing values. The number of sites column indicate this number. Where applicable, we also filtered out sites with MAF less than 0.05 (number shown in next column). All computations were performed in a cluster with 20 cores and 512 GB memory running Red Hat Enterprise Linux (RHEL) 8 operating system.

## 3 Results

Among the statistics implemented by `selscan v2.1`, iHS, nSL, and iHH12 use biallelic genotypic data to pinpoint potential regions of recent or ongoing positive selection within genomes of a single population. XP-EHH and XP-nSL do so across populations. For testing performance of phased iHS and XP-EHH computation, we tested against three state-of-the-art tools: `rehh2, hapbin` and `selscan`. For testing unphased statistics, we used `rehh2` and `selscan`. We used the default parameters for `selscan` in all tools (EHH cutoff 0.05, minimum maf of 0.05 and maximum extend to 1 million bp.).

### 3.1 Performance improvement on phased iHS

Table 2 shows benchmarking results on all datasets for computing phased iHS. This is the key result, as the algorithmic efficiency of `selscan v2.1` is better showcased via comparison to `hapbin`, and the only single population multithreaded statistics implemented in `hapbin` is iHS. From Table 2, we see that the best gain over `hapbin` is obtained on chromosome 10, where `selscan v2.1` is 5.3 times faster with 1 thread and 2 times faster with 16 threads. We noticed this general trend of having the speedup against `hapbin` reduced with increasing number of threads; see Section 3.6 more details.

**Table 2:** Time (in hh:mm format) and memory usage (in GB) for iHS computation. Bold entries indicate best performance. * indicates runs that did not finish within 48 hours.

| Threads | Tool | Chr 19 |  | Chr 10 |  | Chr 6 |  | Chr 1 |  | Simulated sweep |  | Simulated neutral |  |
| --- | --- | --- | --- | --- | --- | --- | --- | --- | --- | --- | --- | --- | --- |
|  |  | Time | GB | Time | GB | Time | GB | Time | GB | Time | GB | Time | GB |
| 1 | selscan v2.1 | **0:48** | **2** | **1:50** | **2.8** | **4:12** | **0.7** | **2:51** | **0.8** | **1:23** | **0.2** | **12:37** | 3.7 |
|  | hapbin | 3:46 | **1.5** | 9:43 | 3.2 | 6:31 | 3.6 | 11:44 | 4.4 | 20:10 | 24 | 21:26 | **1.9** |
|  | selscan v2.0 | 38:37 | 10.6 | * | * | * | * | * | * | * | * | * | * |
|  | rehh2 | 5:16 | 275 | * | * | * | * | * | * | * | * | * | * |
| 16 | selscan v2.1 | **0:09** | **2** | **0:13** | **0.7** | **0:20** | **1** | **0:20** | **1** | **1:04** | **0.2** | **1:16** | 3.7 |
|  | hapbin | 0:20 | **1.7** | 0:34 | 2.8 | 0:48 | 3.5 | 0:53 | 4.4 | 6:11 | 28 | 1:30 | **2.5** |
|  | selscan v2.0 | 4:49 | 10.6 | 6:52 | 19.7 | 13:09 | 24.8 | 9:37 | 31.7 | * | * | * | * |
|  | rehh2 | 3:00 | 275 | 6:29 | 453 | 8:32 | 569 | 11:22 | 734 | * | * | * | * |

Generally, for all datasets, runtime is substantially better than both `selscan` and `rehh2`. For 16 threads, we see massive improvement over `rehh2` (chr 19: 20x faster and 138x less memory, chr 1: 34x faster and 733x less memory). On the simulated large neutral dataset, `selscan v2.1` is 49x faster and uses 6x less memory than `selscan` (62hr, 22GB). This result is omitted from Table 2 for consistency, as we exclude runs exceeding 48 hours. Only for this dataset, which required extensive filtering to reduce over 1 billion sites to around 1 million by removing low-MAF variants, we pre-filtered using `vcftools` to ensure a fair comparison. Interestingly, with pre-filtering, `selscan v2.1` achieves only a 1.2x speedup over `hapbin` on 16 threads (as shown in the table). However, when allowing `hapbin` and `selscan v2.1` to perform filtering internally, we observe up to a 6x speedup. We report the pre-filtered results here for fairness.

In some datasets, we see that `hapbin` has slightly better memory usage than `selscan v2.1`, although the difference is small, generally less than 2x.

### 3.2 Performance improvement using multi-parameter configuration

A key advantage of `selscan v2.1` is introduction of multiple parameter support. For large VCF files, time to load data into memory and preparing it for EHH-based statistics calculation is a big overhead. In some datasets we see more than 50% time spent in loading the input genotype data. Therefore, support for calculation of multiple statistics along with varying parameters (i.e. minimum allele frequency cutoff, EHH decay cutoff, maximum number of SNPs to extend etc.) offers major speedup advantage.

We test this on chr1 from 1000 genomes project (Table 3). We run calculations with a minimum MAF cutoff set to 0.01, 0.05, 0.1, 0.15, 0.2 and EHH cutoff set to 0.05 and 0.1. For these 10 sets of parameters, `selscan` and `hapbin` both were run sequentially whereas `selscan v2.1` utilized its multi-parameter support for speedup. Loading the data for each run takes about half an hour, totaling 5 hours for 10 runs, which increases with more parameter sets. Coupled with `selscan v2.1`’s efficient algorithm, this achieves 11x speedup over `hapbin` and uses 2.5x less memory.

**Table 3:** Time and memory for multi-parameter iHS calculation (phased). Number of threads used here is 16.

|  | Time (h) | Memory (gb) |
| --- | --- | --- |
| selscan v2.1 | 2.0 | 1.8 |
| hapbin | 22.5 | 4.6 |
| selscan v2.0 | >24 | 5.5 |

### 3.3 Performance improvement on unphased statistics calculation

We notice in Table 4 that runtime is cut down in half for unphased analysis for `selscan` and `rehh2` . However in `selscan v2.1`, this we see that this improvement is less than two-fold due to the complexity of the unphased algorithm. However, we still see speedup of 3.7x and 18x and memory improvement of 1.6x and 74x over `selscan` and `rehh2` respectively.

**Table 4:** Time and memory usage for phased and unphased iHS using 16 threads.

|  | Unphased iHS |  |  | Phased iHS |  |  |
| --- | --- | --- | --- | --- | --- | --- |
|  | selscan v2.1 | selscan v2.0 | rehh2 | selscan v2.1 | selscan v2.0 | rehh2 |
| Time | 0 h 19 min | 1 h 11 min | 5 h 42 min | 0 h 9 min | 2 h 40 min | 10 h 54 min |
| Memory | 3.7 GB | 5.8 GB | 273.8 GB | 2.0 GB | 11.4 GB | 273.8 GB |

### 3.4 Performance improvement on XP-EHH calculation

The power of XP-statistics is better without MAF filtering. Hence, we tested with the complete dataset for `selscan v2.1, selscan` and `rehh2` (Table 5). As expected when the number of sites is very high, `selscan` and `rehh2` do not scale well.

**Table 5:** Time and memory usage for XP-EHH on chromosome 1 (CHB vs CEU, 4.8M SNPs) without MAF cutoff.

|  | selscan v2.1 | hapbin | selscan v2.0 | rehh2 |
| --- | --- | --- | --- | --- |
| Time | 7 min | 7 min | 13 h 7 min | 14 h 54 min |
| Memory | 7.1 GB | 3.1 GB | 2.9 GB | 35.1 GB |

`selscan v2.1` uses 5 times less memory than `rehh2` . The runtime is 112x and 120x less than `selscan` and rehh2, respectively. We notice that for XP statistics, `selscan v2.1` generally uses slightly more memory than `selscan v2.0` and `hapbin` at higher thread counts, likely due to minor overlooked optimizations during development.

### 3.5 Performance improvement over hapbin on very large sample sizes

Although in Table 2 and Table 5, we see that in most practical datasets, performance of `selscan v2.1` matches or outperforms `hapbin`, there are certain cases where `selscan v2.1` is expected to dramatically outperform `hapbin` . We see that `selscan v2.1` significantly outperforms `hapbin` when the number of sample sizes grows beyond a certain point. In Table 6 and Figure 3, we see that as the number of haplotypes grows from 10K to 500K, `hapbin` performance starts to degrade. We can see a speedup gain of upto 13x. We used `msprime` neutral simulation with the same parameters listed in Table 1, but varied the number of sample sizes to be 10K, 50K, 100K and 500K. We also filtered out sites with MAF < 0.05.

**Table 6:** Speedup and memory improvement of `selscan v2.1` against `hapbin` for large sample sizes.

| Number of Sites ($MAF \geq 0.05$) | Number of Haplotypes | Time (min) | | Memory (GB) | | Time Speedup | Memory Improved by |
| --- | --- | --- | --- | --- | --- | --- | --- |
|  |  | selscan v2.1 | hapbin | selscan v2.1 | hapbin |  |  |
| 12,080 | 10,000 | 1.1 | 1.2 | 0.05 | 0.4 | 1 | 7 |
| 11,564 | 50,000 | 4 | 24 | 0.16 | 7.3 | 6 | 46 |
| 11,530 | 100,000 | 6 | 48 | 0.31 | 8.0 | 8 | 26 |
| 11,888 | 500,000 | 33 | 362 | 1.52 | 15.59 | 11 | 10 |
| 11,604 | 1,000,000 | 58 | 770 | 2.96 | 33.89 | 13 | 11 |

**Figure 3:**
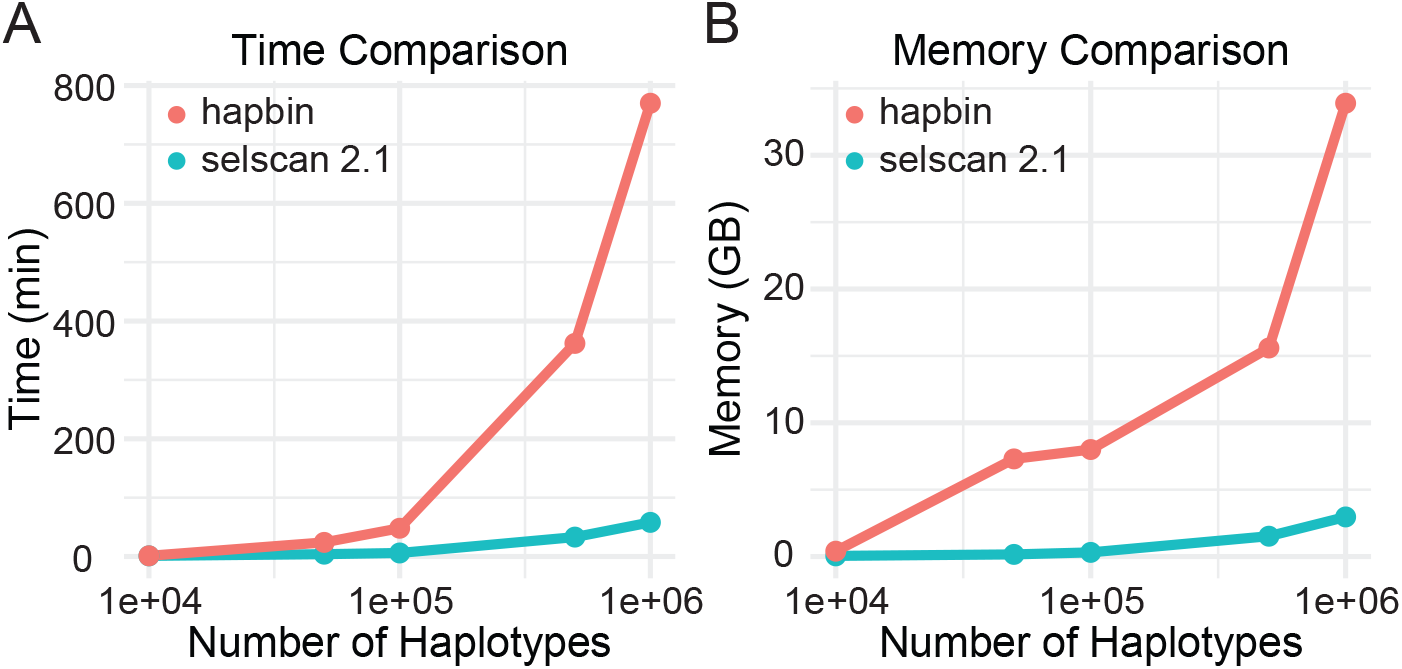
Scaling of time and memory with increasing number of individuals for simulated neutral dataset using 16 threads for `selscan v2.1` (blue line) and `hapbin` (red line). The number of filtered SNPs are around 11K-12K. It shows that although performance of `hapbin` and `selscan v2.1` are similar for sample sizes of 10K or fewer, as it grows beyond that, `selscan v2.1` begins to significantly outperform `hapbin` (data shown in Table 6).

### 3.6 Implications of great performance gain in single-threaded execution

We observe greater speedup in the single-threaded version compared to the multi-threaded one. Specifically, `selscan v2.1` achieves the best runtime across all tools and datasets tested in single-threaded mode. This is likely due to improvements in the core algorithm and data structures, which make the base computation significantly faster. The lower relative speedup with multiple threads occurs because multithreading is most beneficial when there is a large workload to divide. When the overall workload is smaller, there is less opportunity for parallel gains.

Running faster on a single core offers many advantages. It reduces resource usage, which is especially valuable when computing power is limited or costly. It also enables more tasks to be executed simultaneously. Our optimized single-threaded implementation ensures strong performance even on modest hardware, making the tool broadly accessible and efficient.

### 3.7 Validating accuracy of results

We validated all our results by computing correlation with `selscan v2.0` . All parameters are handled the same way as in `selscan v2.0`, only the core EHH computation algorithm is improved. Therefore, we observe a correlation of 0.99 to 1 with `selscan` results for the unstandardized statistics. The minor numerical differences are due to slight variations in how floating-point sums and rounding are handled. The recurrence relationship in Eq. 5 along with its proof in supplementary demonstrates that EHH-based computations are identical to those in `selscan v2.0` . Both `selscan v2.0` and `selscan v2.1` compute exactly the same statistic, with minor numerical deviations arising only from floating-point boundary conditions. Aside from these negligible effects, results show near-perfect correlation with `selscan v2.0` . For all results reported in this paper, we verified that the correlation with `selscan v2.0` is 1.0.

Since `rehh` and `hapbin` do not implement other statistics in same manner, we verify agreement using iHS results. We focus on iHS in particular, as it shares the same settings and provides a direct comparison. We can compare the scores on the simulated dataset across different genomic positions to verify they have same values when standardized by allele frequency bins. For `hapbin` and `rehh2`, all correlation values for values reported in Table 2 were over 0.96.

We performed a thorough simulation study to explicitly demonstrate that the new implementation does not miss selective events or deviate from the ground truth. Specifically, we simulated a 2.5 Mb segment with a selective sweep at the center (selection coefficient *s* = 0.1, recombination rate 1.29 *×* 10^*−*8^, mutation rate 1*×*10^*−*8^), using `SLiM` followed by recapitation in `msprime`. For the cross-population simulation, we introduced a split 500 generations prior to the sweep so that the selected allele segregates at ∼ 70–80% frequency, a regime where detection is typically most informative by these statistics. We then ran `selscan v2.1` on these simulated datasets to compute all statistics. All values were normalized using 100 allele-frequency bins to remove the dependency of iHS/nSL/XP-EHH/iHH12 on allele frequency, ensuring that extreme values reflect true signals of selection rather than frequency-related artifacts. For smoother visualization, values were averaged across 50kb windows. In Figure 4A, the plots of iHS show strong agreement with results from other tools across different chromosomal regions. Across all statistics, the sweep was correctly identified by `selscan v2.1`, as we can see from the intense scores in the middle (Figure 4B,C). These results confirm that runtime improvements do not compromise accuracy.

**Figure 4:**
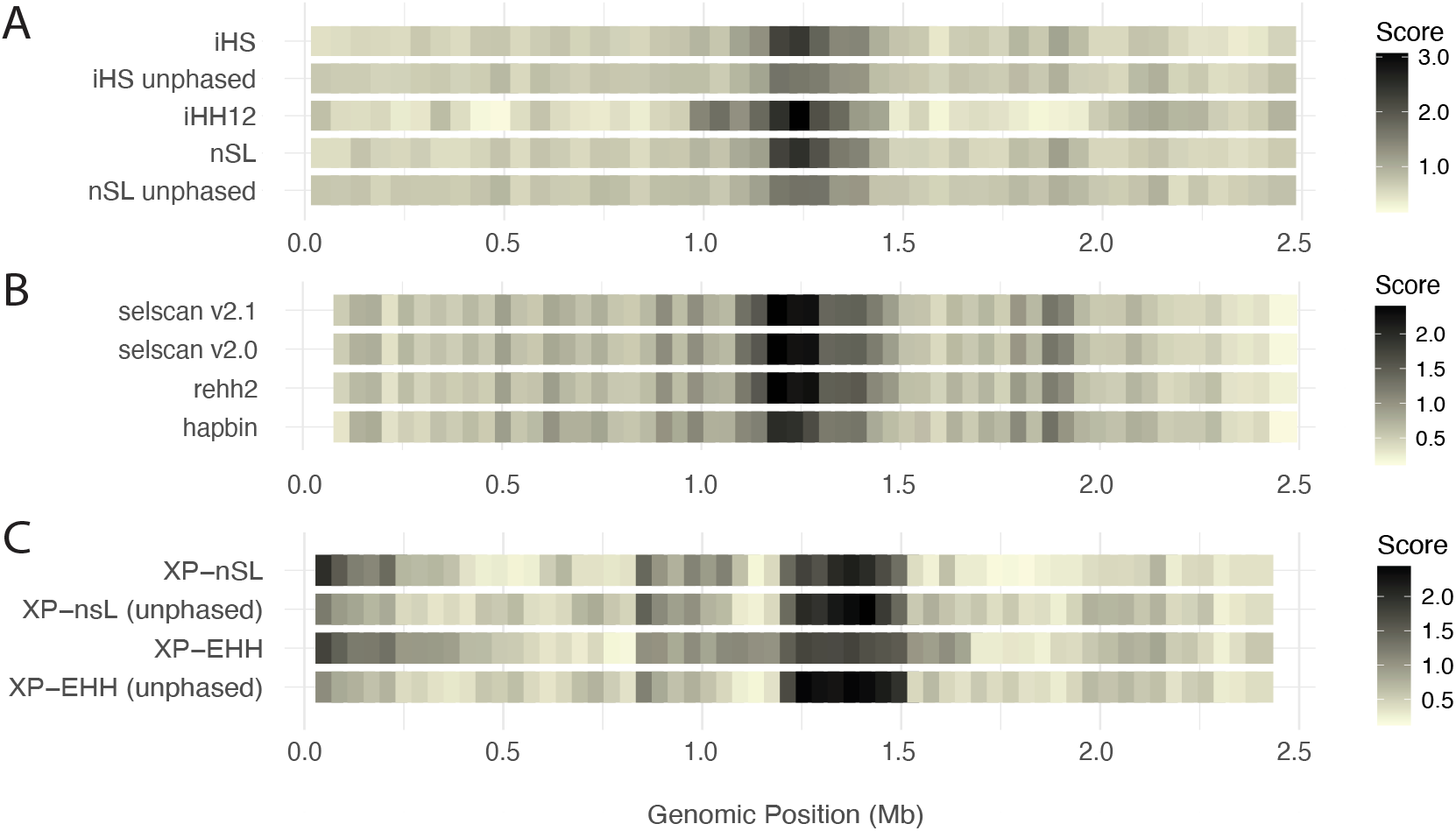
Heatmap of scores across genomic regions for detecting selective sweeps in a simulated experiment. (A) Normalized iHS scores computed with different tools. (B) Single population statistics computed with `selscan v2.1`. (C) Two population statistics computed with `selscan v2.1` . Scores are normalized within frequency bins, and all statistics show the center with the highest values.

### 3.8 Other statistics

The details of the correlation, runtime and memory usage analysis for nSL, XP-nsl, iHH12 are in supplementary (Table S1).

### 3.9 Time and memory improvement as a result of optimizations

Memory usage of `selscan v2.1` is always smaller compared to `rehh2. selscan v2.0` and `hapbin` both have small memory footprints for small datasets, so the difference with `selscan v2.1` is relatively small. However, `rehh2` consumes substantially higher memory than all the other tools. In order to highlight the time and memory tradeoff between tools, we draw a comparison using Chr 19 as an example to visualize this tradeoff (Figure 5). Despite differences in memory and runtime, the tools report highly consistent results as shown in Table 7.

**Table 7:** Pearson correlation between unstandardized iHS values for Chr 19 reported by all tools tested (lower triangle only).

|  | selscan v2.1 | selscan v2.0 | rehh2 | hapbin |
| --- | --- | --- | --- | --- |
| selscan v2.0 | 1.000 | – | – | – |
| rehh2 | 0.996 | 0.996 | – | – |
| hapbin | 0.965 | 0.965 | 0.970 | – |

**Figure 5:**
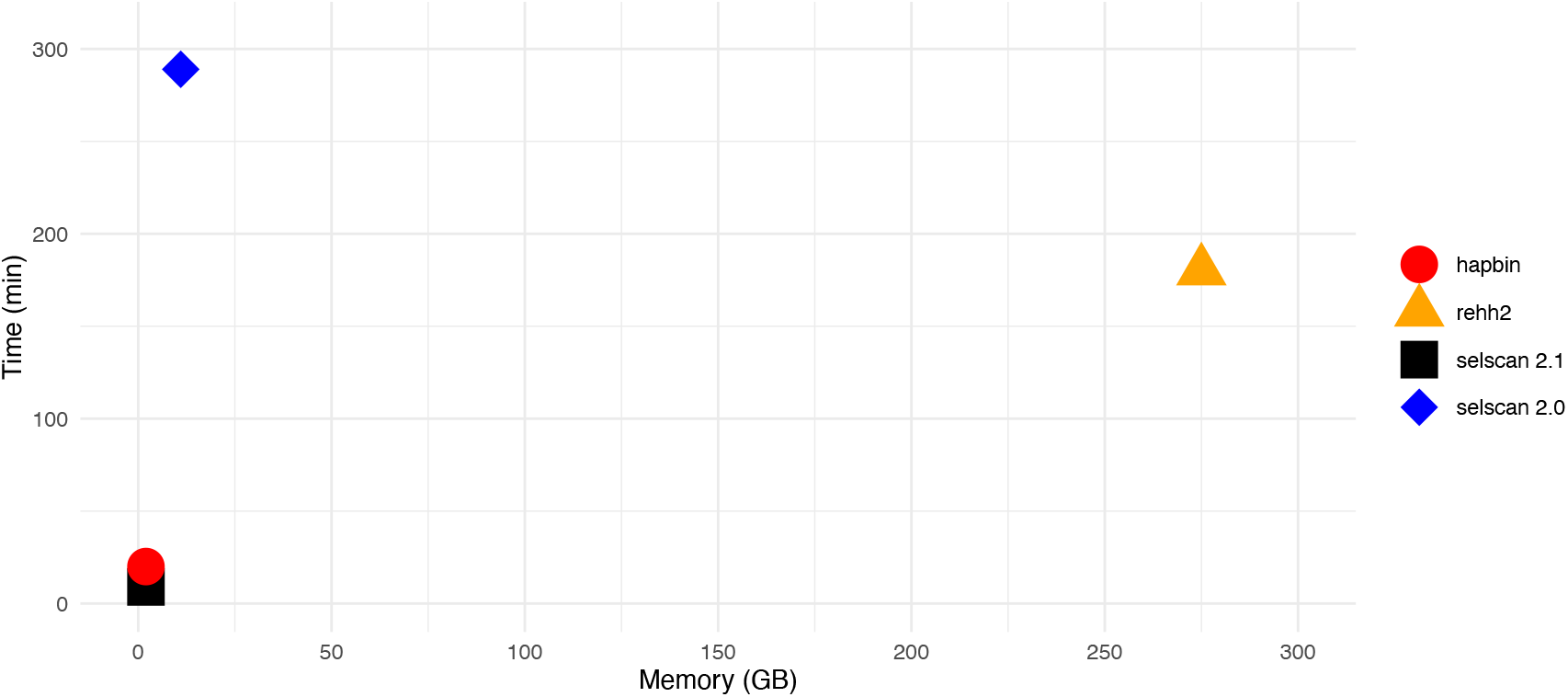
Time and memory for all tools for Chr 19 using 16 threads. Notice that both `hapbin` and `selscan v2.1` consume very little memory and time. Between `selscan` (26x less memory than `rehh2` ) and `rehh2` (1.6x less time than `selscan` ), there is a memory-time tradeoff.

For large datasets, `selscan v2.1` uses dramatically less memory (Figure 3B). We always see runtime improvement in the range of 20x-50x compared to `selscan v2.0` and 4x-34x compared to `rehh2`. This is only based on the results presented in this paper, and we expect to see more improvements in datasets of larger scale, especially if number of sites is increased.

The major performance improvement came from two factors: the use of bitsets and iterating over set bits when adding a new site, rather than relying on haplotype counting via hashing. This is crucial to reduce both runtime and memory usage. Modern compilers enable linear-time iteration over a bitset relative to the number of set bits (not the bitset’s total size) by using efficient bitwise operations and hardware instructions for counting trailing zeroes to locate the next set bit. This allows for the memory usage gain with no additional runtime cost. Multithreading also improves performance, and it is optimized in selscan v2.1 with the use of a thread pool. However even with a thread pool, because the time to compute EHH largely depends on LD decay patterns around the particular locus, the workload is not always evenly distributed among all threads.

## 4 Discussion

We implemented fast and memory efficient versions of several popular EHH-based selection statistics, including both phased and unphased implementations. Because the main strength of the newly proposed tool is its scalability, we mostly chose our datasets to have large number of variants so that its advantage over other tools become apparent. To maintain good power for detecting the sweeps, EHH-based statistics need to be computed over a large genomic region, so scalability of these computations is critically important. When customizing parameters for different model and non-model organisms, the results presented here offer reassurance that the process can be completed within a reasonable timeframe.

We have shown through extensive experiments that `selscan v2.1` substantially optimizes these computations. A notable use case is rapid generation of large-scale multi-parameter training dataset for machine learning models to detect selective sweeps. While other tools could theoretically generate datasets of this scale, they would require order of magnitude more time to complete the task as shown by our experiments. Another key advantage of `selscan v2.1` lies in its efficient memory usage. Compared to the widely used `rehh2, selscan v2.1` achieves substantial memory reductions, making it possible to perform large-scale analyses even on personal computers.

Both `hapbin` and `selscan v2.1` utilize bitsets to achieve a lower memory footprint; however, their algorithms differ significantly. `Hapbin` maintains a tree of bitsets in each iteration, which can theoretically exceed O(K) complexity and scale with the increasing number of haplotypes and SNPs. Although we did not encounter this scalability issue in our 1000-genome dataset, simulations reveal this limitation. From simulation, we see significant improvement over `hapbin` in handling larger populations (Table 6 and Figure 3). We note that `selscan v2.1` does still outperform `hapbin` in computing iHS for datasets where the number of samples and variants is small, although runtimes are generally short in these contexts. However, when generating training data for ML approaches, even small improvements in runtime can add up substantially (Table 3). In general, our improved algorithm for computing EHH-based selection statistics offers better runtime and memory efficiency along with improved customization options over `hapbin, selscan v2.0`, and `rehh2`. These improvements are particularly notable for very large datasets. `selscan v2.1` is available at https://github.com/szpiech/selscan.

## 5 Acknowledgments

Computations for this research were performed using the Pennsylvania State University’s Institute for Computational Data Sciences’ Roar supercomputer. This work was supported by the National Institute of General Medical Sciences of the National Institutes of Health award number R35GM146926, and start-up funds from the Pennsylvania State University’s Department of Biology.

## References

Alonso-Blanco, C., Andrade, J., Becker, C., Bemm, F., Bergelson, J., Borgwardt, K. M., Cao, J., Chae, E., Dezwaan, T. M., Ding, W., et al. (2016). 1,135 genomes reveal the global pattern of polymorphism in arabidopsis thaliana. Cell, 166(2):481–491.

Arnab, S. P., Amin, M. R., and DeGiorgio, M. (2023). Uncovering footprints of natural selection through spectral analysis of genomic summary statistics. Molecular Biology and Evolution, 40(7):msad157.

Bycroft, C., Freeman, C., Petkova, D., Band, G., Elliott, L. T., Sharp, K., Motyer, A., Vukcevic, D., Delaneau, O., O’Connell, J., et al. (2018). The uk biobank resource with deep phenotyping and genomic data. Nature, 562(7726):203–209.

Cardona, A., Pagani, L., Antao, T., Lawson, D. J., Eichstaedt, C. A., Yngvadottir, B., Shwe, M. T. T., Wee, J., Romero, I. G., Raj, S., et al. (2014). Genome-wide analysis of cold adaptation in indigenous siberian populations. PloS one, 9(5):e98076.

DeGiorgio, M. and Szpiech, Z. A. (2022). A spatially aware likelihood test to detect sweeps from haplotype distributions. PLoS genetics, 18(4):e1010134.

Durbin, R. (2014). Efficient haplotype matching and storage using the positional burrows–wheeler transform (pbwt). Bioinformatics, 30(9):1266–1272.

Ferrer-Admetlla, A., Liang, M., Korneliussen, T., and Nielsen, R. (2014). On detecting incomplete soft or hard selective sweeps using haplotype structure. Molecular biology and evolution, 31(5):1275–1291.

Garrison, E., Kronenberg, Z. N., Dawson, E. T., Pedersen, B. S., and Prins, P. (2021). Vcflib and tools for processing the vcf variant call format. BioRxiv, 21.

Garrison, E., Kronenberg, Z. N., Dawson, E. T., Pedersen, B. S., and Prins, P. (2022). A spectrum of free software tools for processing the vcf variant call format: vcflib, bio-vcf, cyvcf2, hts-nim and slivar. PLoS Computational Biology, 18(5):e1009123.

Gautier, M., Klassmann, A., and Vitalis, R. (2017). rehh 2.0: a reimplementation of the r package rehh to detect positive selection from haplotype structure. Molecular ecology resources, 17(1):78– 90.

Gautier, M. and Vitalis, R. (2012). rehh: an r package to detect footprints of selection in genomewide snp data from haplotype structure. Bioinformatics, 28(8):1176–1177.

Harris, A. M. and DeGiorgio, M. (2020). A likelihood approach for uncovering selective sweep signatures from haplotype data. Molecular biology and evolution, 37(10):3023–3046.

Harris, A. M., Garud, N. R., and DeGiorgio, M. (2018). Detection and classification of hard and soft sweeps from unphased genotypes by multilocus genotype identity. Genetics, 210(4):1429–1452.

Hartmann, F. E., Vonlanthen, T., Singh, N. K., McDonald, M. C., Milgate, A., and Croll, D. (2021). The complex genomic basis of rapid convergent adaptation to pesticides across continents in a fungal plant pathogen. Molecular Ecology, 30(21):5390–5405.

Hawkins, N. J., Bass, C., Dixon, A., and Neve, P. (2019). The evolutionary origins of pesticide resistance. Biological Reviews, 94(1):135–155.

Hejase, H. A., Dukler, N., and Siepel, A. (2020). From summary statistics to gene trees: methods for inferring positive selection. Trends in Genetics, 36(4):243–258.

Hemstrom, W., Grummer, J. A., Luikart, G., and Christie, M. R. (2024). Next-generation data filtering in the genomics era. Nature Reviews Genetics, pages 1–18.

Ingram, C. J., Mulcare, C. A., Itan, Y., Thomas, M. G., and Swallow, D. M. (2009). Lactose digestion and the evolutionary genetics of lactase persistence. Human genetics, 124:579–591.

Kelleher, J., Etheridge, A. M., and McVean, G. (2016). Efficient coalescent simulation and genealogical analysis for large sample sizes. PLoS computational biology, 12(5):e1004842.

Kern, A. D. and Schrider, D. R. (2018). diplos/hic: an updated approach to classifying selective sweeps. G3: Genes, Genomes, Genetics, 8(6):1959–1970.

Lauterbur, M. E., Munch, K., and Enard, D. (2023). Versatile detection of diverse selective sweeps with flex-sweep. Molecular Biology and Evolution, 40(6):msad139.

Lin, K., Li, H., Schlotterer, C., and Futschik, A. (2011). Distinguishing positive selection from neutral evolution: boosting the performance of summary statistics. Genetics, 187(1):229–244.

Liu, X., Zhang, Y., Li, Y., Pan, J., Wang, D., Chen, W., Zheng, Z., He, X., Zhao, Q., Pu, Y., et al. (2019). Epas1 gain-of-function mutation contributes to high-altitude adaptation in tibetan horses. Molecular biology and evolution, 36(11):2591–2603.

Lowy-Gallego, E., Fairley, S., Zheng-Bradley, X., Ruffier, M., Clarke, L., Flicek, P., Consortium, . G. P., et al. (2019). Variant calling on the grch38 assembly with the data from phase three of the 1000 genomes project. Wellcome open research, 4.

Maclean, C. A., Chue Hong, N. P., and Prendergast, J. G. (2015). Hapbin: an efficient program for performing haplotype-based scans for positive selection in large genomic datasets. Molecular biology and evolution, 32(11):3027–3029.

Melsted, P. and Pritchard, J. K. (2011). Efficient counting of k-mers in dna sequences using a bloom filter. BMC bioinformatics, 12:1–7.

Miles, A., pyup.io bot, Rodrigues, M. F., Ralph, P., Kelleher, J., Schelker, M., Pisupati, R., Rae, S., and Millar, T. (2024). cggh/scikit-allel: v1.3.13.

Nawaz, M. Y., Savegnago, R. P., Lim, D., Lee, S. H., and Gondro, C. (2024). Signatures of selection in angus and hanwoo beef cattle using imputed whole genome sequence data. Frontiers in Genetics, 15:1368710.

Pybus, M., Luisi, P., Dall’Olio, G. M., Uzkudun, M., Laayouni, H., Bertranpetit, J., and Engelken, J. (2015). Hierarchical boosting: a machine-learning framework to detect and classify hard selective sweeps in human populations. Bioinformatics, 31(24):3946–3952.

Sabeti, P. C., Varilly, P., Fry, B., Lohmueller, J., Hostetter, E., Cotsapas, C., Xie, X., Byrne, E. H., McCarroll, S. A., Gaudet, R., et al. (2007). Genome-wide detection and characterization of positive selection in human populations. Nature, 449(7164):913–918.

Spence, J. P., Zeng, T., Mostafavi, H., and Pritchard, J. K. (2023). Scaling the discrete-time wright–fisher model to biobank-scale datasets. Genetics, 225(3):iyad168.

Sugden, L. A., Atkinson, E. G., Fischer, A. P., Rong, S., Henn, B. M., and Ramachandran, S. (2018). Localization of adaptive variants in human genomes using averaged one-dependence estimation. Nature communications, 9(1):703.

Szpiech, Z. A. (2024). selscan 2.0: scanning for sweeps in unphased data. Bioinformatics, 40(1):btae006.

Szpiech, Z. A. and Hernandez, R. D. (2014). selscan: an efficient multithreaded program to perform ehh-based scans for positive selection. Molecular biology and evolution, 31(10):2824–2827.

Szpiech, Z. A., Novak, T. E., Bailey, N. P., and Stevison, L. S. (2021). Application of a novel haplotype-based scan for local adaptation to study high-altitude adaptation in rhesus macaques. Evolution letters, 5(4):408–421.

Taliun, D., Harris, D. N., Kessler, M. D., Carlson, J., Szpiech, Z. A., Torres, R., Taliun, S. A. G., Corvelo, A., Gogarten, S. M., Kang, H. M., et al. (2021). Sequencing of 53,831 diverse genomes from the nhlbi topmed program. Nature, 590(7845):290–299.

Tishkoff, S. A., Reed, F. A., Ranciaro, A., Voight, B. F., Babbitt, C. C., Silverman, J. S., Powell, K., Mortensen, H. M., Hirbo, J. B., Osman, M., et al. (2007). Convergent adaptation of human lactase persistence in africa and europe. Nature genetics, 39(1):31–40.

Vatsiou, A. I., Bazin, E., and Gaggiotti, O. E. (2016). Detection of selective sweeps in structured populations: a comparison of recent methods. Molecular ecology, 25(1):89–103.

Voight, B. F., Kudaravalli, S., Wen, X., and Pritchard, J. K. (2006). A map of recent positive selection in the human genome. PLoS biology, 4(3):e72.

Xue, A. T., Schrider, D. R., and Kern, A. D. (2021). Discovery of ongoing selective sweeps within anopheles mosquito populations using deep learning. Molecular biology and evolution, 38(3):1168–1183.

Zhao, S., Chi, L., Fu, M., and Chen, H. (2024). Haplosweep: detecting and distinguishing recent soft and hard selective sweeps through haplotype structure. Molecular Biology and Evolution, 41(10):msae192.

