## Supplementary Materials for "Fast and Memory-Efficient Dynamic Programming Approach for Large-Scale EHH-Based Selection Scans"

#### S1 Proof of the EHH Recurrence for Efficient Computation

We recall the notation $\mathcal{H}_m$ from the main text. It denotes all the $m$ length haplotypes extending to $m$.

$$EHH(m) = \sum_{x \in \mathrm{Supp}(\mathcal{H}_m)} \frac{\binom{m_{\mathcal{H}_m}(x)}{2}}{\binom{K}{2}} \tag{S1}$$

We want to prove that instead of Eq. S1 we can replace with the following relations Eq. S2 and Eq. S3.

$$EHH(m) = EHH(m-1) - \delta_m \tag{S2}$$

$$\delta_m = \frac{1}{\binom{K}{2}} \sum_{h \in \mathrm{Supp}(\mathcal{H}_{m-1})} m_{\mathcal{H}_m}(h \odot 0) \cdot m_{\mathcal{H}_m}(h \odot 1) \tag{S3}$$

**Lemma S1.1.** *[Family counts and splitting] For each prefix $h \in \mathrm{Supp}(\mathcal{H}_{m-1})$, define*

$$n_0^{(h)} := m_{\mathcal{H}_{m-1}}(h \odot 0), \quad n_1^{(h)} := m_{\mathcal{H}_{m-1}}(h \odot 1),$$

*Then,*

$$n^{(h)} := n_0^{(h)} + n_1^{(h)} = m_{\mathcal{H}_m}(h).$$

*We say that the family $h$* splits *at locus $m$ if $n_0^{(h)} \geq 1$ and $n_1^{(h)} \geq 1$, and denote $\mathcal{G}_{01} := \{h \in \mathrm{Supp}(\mathcal{H}_{m-1}) : n_0^{(h)} \geq 1$ and $n_1^{(h)} \geq 1\}$.*

**Theorem S1.2** (Recurrence Relation for EHH)**.** *For every $m > 0$,*

$$EHH(m) = EHH(m-1) - \delta_m$$

*where*

$$\delta_m = \frac{1}{\binom{K}{2}} \sum_{h \in \mathrm{Supp}(\mathcal{H}_{m-1})} n_0^{(h)} n_1^{(h)}$$

*and*

$$EHH(0) = 1$$

*Proof.* We proceed by induction. The base case is satisfied immediately. The multiplicity for the core locus ($m = 0$) is $K$, and therefore, $EHH(0) = 1$. Now, we assume that the equation holds for a positive integer $m - 1$. We wish to show that the result extends to the case of $m$. We begin with

$$EHH(m) = \sum_{x \in \mathrm{Supp}(\mathcal{H}_m)} \frac{\binom{m_{\mathcal{H}_m}(x)}{2}}{\binom{K}{2}}$$

By Eq. S1, we can group the pairs according to their $(m-1)$-length prefix $h \in \mathrm{Supp}(\mathcal{H}_{m-1})$:

$$EHH(m) = \frac{1}{\binom{K}{2}} \sum_{h \in \mathrm{Supp}(\mathcal{H}_{m-1})} \left( \binom{n_0^{(h)}}{2} + \binom{n_1^{(h)}}{2} \right)$$

Using the binomial identity $\binom{a}{2} + \binom{b}{2} = \binom{a+b}{2} - ab$ with $a = n_0^{(h)}, b = n_1^{(h)}$. The total number of pairs is the sum of the pairs within the first group, the pairs within the second group, and the pairs that include one element from each group. We have

$$EHH(m) = \frac{1}{\binom{K}{2}} \sum_{h \in \text{Supp}(\mathcal{H}_{m-1})} \binom{n_0^{(h)} + n_1^{(h)}}{2} - n_0^{(h)} n_1^{(h)}.$$

By our induction hypothesis and Eq. S1,

$$EHH(m) = E(m-1) - \frac{1}{\binom{K}{2}} \sum_{h \in \text{Supp}(\mathcal{H}_{m-1})} n_0^{(h)} n_1^{(h)}.$$

We note that each term $n_0^{(h)} n_1^{(h)} \geq 0$, so $EHH(m)$ is non-increasing in $m$. This completes the proof. □

**Corollary S1.3** (Summing only over splitting families)**.** *The decrement can be restricted to splitting families:*

$$\delta_m = \frac{1}{\binom{K}{2}} \sum_{h \in \mathcal{G}_{01}} n_0^{(h)} n_1^{(h)}.$$

*Proof.* If a family $h$ does not split at locus $m$, then either $n_0^{(h)} = 0$ or $n_1^{(h)} = 0$, so its contribution $n_0^{(h)} n_1^{(h)} = 0$. Hence, restricting the sum to $\mathcal{G}_{01}$ does not change its value. □

**Proposition S1.4** (Work bound per step)**.** *Let $p_m$ be the number of haplotypes carrying allele 1 at locus $m$. Then*

$$\#\mathcal{G}_{01} \leq \min\{p_m, K - p_m\} \leq p_m,$$

*and computing $\delta_j$ by Definition S1.3 requires $\Theta(\#\mathcal{G}_{01}) = \mathcal{O}(\min\{p_m, K - p_m\}) \subseteq \mathcal{O}(p_m)$ time.*

*Proof.* Each splitting family has at least one haplotype with allele 1 and one with allele 0. Thus, the number of splitting families is at most the number of haplotypes with allele 1,

$$\#\mathcal{G}_{01} \leq p_m,$$

and also at most the number with allele 0,

$$\#\mathcal{G}_{01} \leq K - p_m.$$

The sum in Eq. S1 runs over exactly these splitting families, giving the stated time bound. □

#### S2 Additional results (iHH12, nSL, and XP-nSL)

For iHH12, nSL, and XP-nSL, we did benchmarking only against `selscan v2.0`, as other tools did not implement these statistics. The results were validated using correlation tests. `Selscan v2.1` shows substantial performance improvements over `selscan v2.0` in computing ihh12, nSL, and XP-nSL in chr1 datasets. For XP-nSL, CEU and CHB populations as referenced in main text are used. In single-threaded scenarios, speedups range from approximately 15x to over 50x. When using 16 threads, `selscan v2.1` achieves speedups between 6.7x and 21.8x. Note that, for the XP-nSL, `selscan v2.1` exhibits slightly higher memory usage at higher thread counts. We believe it may have been due to certain memory optimizations that were inadvertently overlooked for XP-statistics during development.

Table S1: Performance comparison of `selscan v2.1` and `selscan v2.0` for different thread configurations on chr1.

| n. Threads | Tool | iHH12 | | nSL | | XP-nSL | |
|---|---|---|---|---|---|---|---|
| | | Time (h) | Memory (GB) | Time (h) | Memory (GB) | Time (h) | Memory (GB) |
| | selscan v2.1 | 0 hr 27 min | 0.9 | 0 hr 26 min | 0.9 | 0 hr 5 min | 4.3 |
| 16 | selscan v2.0 | 3 hr 55 min | 31.7 | 2 hr 55 min | 31.7 | 1 hr 49 min | 2.8 |
| 1 | selscan v2.1 | 1 hr 55 min | 0.9 | 1 hr 52 min | 0.9 | 0 hr 27 min | 2.7 |
| | selscan v2.0 | 34 hr 19 min | 31.7 | 28 h 1 min | 31.7 | 24 h 37 min | 2.8 |

### S3 Algorithm details

Refer to the pseudocode in Algorithm S1 for algorithm details.

#### S3.1 Detailed pseudocode

**Algorithm S1:** Incremental EHH Update via Haplotype Color Splitting

```
Input: Binary haplotype matrix H of size K × N, focal locus i
Output: Color and count arrays for each locus m, updated EHH(m) values
Initialization at m = 0 (i.e., locus i):
    color_prev[0...K−1] ← 0 ;                    // All haplotypes in one group
    count_prev[0] ← K
    numColors_prev ← 1
    Compute EHH(0) using Section 2.2.1
for m = 1 to U do
    // Initialize current column state
    color_curr[0...K−1] ← −1
    count_curr[0...K−1] ← 0
    colorMap ← empty map ;    // Stores indices of haplotypes with allele 1 grouped by
     previous color
    pair ← empty map
    numColors_curr ← numColors_prev
    // Group haplotypes with 1 at position i + m by their previous color
    for k = 0 to K−1 do
        if H[k][i+m] = 1 then
            c ← color_prev[k]
            Append k to colorMap[c]
    // Process each color group
    foreach color c in colorMap do
        members ← colorMap[c]
        if len(members) < count_prev[c] then
            // Split occurs
            new_c ← numColors_curr
            foreach k in members do
                color_curr[k] ← new_c
            for k = 0 to K−1 do
                if color_prev[k] = c and H[k][i+m] = 0 then
                    color_curr[k] ← c
            pair[c] ← new_c; pair[new_c] ← c
            count_curr[c] ← count_prev[c] − len(members)
            count_curr[new_c] ← len(members)
            numColors_curr ← numColors_curr +1
        else
            for k = 0 to K−1 do
                if color_prev[k] = c then
                    color_curr[k] ← c
            count_curr[c] ← count_prev[c]
    // Update EHH using current counts
    EHH[m] ← EHH[m−1] −δ_m
    Compute δ_m = Σ_c count_curr[c] · count_curr[pair[c]]/(K choose 2)
    // Prepare for next iteration
    color_prev ← color_curr
    count_prev ← count_curr
    numColors_prev ← numColors_curr
```

##### S3.2 Theoretical Runtime and Memory Usage

To compute genome-wide iHS, Eq. 2, Eq. 3, and Eq. 4 are computed for all loci $i = 0$ to $N-1$. Without any cutoff, the upstream limit $U$ would be $N-1$, the length of the chromosome. However, in practice, it is unnecessary to compute $EHH(i, j)$ for distant markers once haplotypes become nearly unique, as the homozygosity decays rapidly. Thus, the maximum extent of EHH integration is determined by:

$$U = \min(\texttt{cutoff}, |N-1-i|, \texttt{maxbp}),$$

where `cutoff` is the locus beyond which EHH falls below a user-defined threshold (e.g., 0.05), and `maxbp` is the maximum allowed base-pair distance from the core.

This formulation enables a direct comparison of the runtime complexity between `selscan v2.0` and `selscan v2.1`. In `selscan v2.0`, computing a single $EHH(i, j)$ requires $O(K \cdot |j-i|)$ time due to repeated string hashing. Summing over all upstream and downstream positions gives an overall runtime of $O(K(U^2 + D^2))$, where $U$ and $D$ are the upstream and downstream integration limits, respectively.

In contrast, `selscan v2.1` leverages the recurrence formulation in Eq. 5, computing each $EHH(i, j)$ in time proportional to the number of 1s at locus $j$, i.e., $\Theta(|p_j|) = O(K)$. The total runtime becomes $\Theta(p) = O(K(U + D))$, where $p = \sum_j |p_j|$ across all $L = U + D$ loci.

Only a minimal set of data structures is maintained in memory during each iteration—specifically, the vectors $\texttt{count}^{j-1}$, $\texttt{color}^{j-1}$, and the scalar $EHH^{j-1}$. This results in a space complexity of $O(K)$ per iteration. In practice, overall memory usage is bounded by the input size.

#### S4 Description of multiple parameters supported by `selscan v2.1`

Several parameters guide the runtime and results of the statistics described. Here, we briefly describe the parameters and explain why using a multi-parameter configuration is useful for improving the robustness of the statistics.

**Using multiple statistics on one input** Sometimes, selection signals are best detected by comparing results from multiple statistics. Instead of running each one separately, it is more efficient to compute them all at once, reducing the heavy input load time.

**MAF Filtering** MAF filtering is important in EHH-based calculations. With a too-stringent cutoff, rare alleles may not form strong haplotypes, making EHH decay unpredictable. On the other hand, including very common variants may dilute selection signals if those variants are not under recent selection.

**Distance from Core Haplotype** The genetic or physical distance (in base pairs or centimorgans) from the core haplotype, beyond which calculation is stopped.

**Core Haplotype Frequency** If the frequency of the core haplotype in the population is below a defined threshold, it is excluded from the analysis.

**Threshold for EHH Calculation** A cutoff point (e.g., where EHH falls below 0.05) that determines how far the extended haplotype is considered to persist.

#### S5 Command lines used

In our experiments, we used default parameters for selscan (v2.0.3) and used the same parameters for hapbin and rehh2 where possible. Default values for selscan are: EHH cutoff 0.05, min-maf 0.05, gap-scale 20,000, max-gap 200,000, max-extend 1,000,000. We set --trunc-ok flag to not discard markers at border. Here we show command lines for chr19 dataset with 16 threads.

##### Selscan v2.0.3 and v2.1

```
selscan --vcf chr19.vcf --ihs --pmap --trunc-ok --threads 16
```

##### REHH v2.0

```
hap <- data2haplohh(chr19.vcf, verbose = TRUE, min_maf=0.05, allele_coding =
↪  "01", vcf_reader="vcfR", polarize_vcf=F, remove_multiple_markers=T)
res.ihh<-scan_hh(hap,discard_integration_at_border = FALSE, threads=16,
↪  scalegap=20000, maxgap=200000)
```

In order to obtain the hap file required for hapbin we use

```
bcftools convert --haplegendsample chr19 --vcf chr19.vcf.gz
```

##### Hapbin

```
module load gcc/13.2.0; export OMP_NUM_THREADS=16;
ihsbin --map chr19.map --hap chr19.hap max-extend 1000000 --binom --scale 20000
```
